# Three-dimensional interrelationship between osteocyte network and forming mineral during human bone remodeling

**DOI:** 10.1101/2020.11.20.391862

**Authors:** Mahdi Ayoubi, Alexander F. van Tol, Richard Weinkamer, Paul Roschger, Peter C. Brugger, Andrea Berzlanovich, Luca Bertinetti, Andreas Roschger, Peter Fratzl

## Abstract

During bone remodeling, osteoblasts are known to deposit unmineralized collagenous tissue (osteoid), which mineralizes after some time lag. Some of the osteoblasts differentiate into osteocytes, forming a cell network within the lacunocanalicular network (LCN) of bone. To get more insight into the potential role of osteocytes in the mineralization process of osteoid, sites of bone formation were three-dimensionally imaged in nine forming human osteons using focused ion beam-scanning electron microscopy (FIB-SEM). In agreement with previous observations, the mineral concentration was found to gradually increase from the central Haversian canal towards preexisting mineralized bone. Most interestingly, a similar feature was discovered on a length scale more than 100-times smaller, whereby mineral concentration increased from the LCN, leaving around the canaliculi a zone virtually free of mineral, the size of which decreases with progressing mineralization. This suggests that the LCN controls mineral formation but not just by diffusion of mineralization precursors, which would lead to a continuous decrease of mineral concentration from the LCN. Our observation is, however, compatible with the co-diffusion and reaction of precursors and inhibitors from the LCN into the bone matrix.

## Introduction

Bone is a remarkable material since it reconciles the otherwise conflicting mechanical properties of being difficult to deform (i.e., high stiffness) and difficult to break (i.e., high toughness)^1, 2, 3, 4^. These functional characteristics are attributed to its structure at the macro-, micro-, and nanoscale ^5, 6^. At the smallest length scale, of particular importance is the incorporation of nanoscopic mineral particles of carbonated hydroxyapatite into a soft collagen matrix, which enables a high energy absorption before failure. The mechanical integrity is further improved by a continuous renewal of bone, which allows the removal of microdamage counteracting the fatigue of the material ^7, 8, 9, 10^. This remodeling of bone occurs in cortical bone of humans and bigger mammals by the formation of osteons. Osteons are roughly cylindrical structures with a diameter of about 300 μm and a central cavity known as the Haversian canal, which provides space for blood vessels and nerves. They are formed by bone-resorbing osteoclasts tunneling through the cortical bone and in their wake, osteoblasts lay down new bone, thereby narrowing down the tunnel until only the Haversian canal remains. Such a replacement of old by new bone is very different to bone formation during embryonic development, which occurs via the intermediate stage of cartilage in a process called endochondral ossification. Furthermore, the remodeling scenario of a forming osteon has to be distinguished from modeling, where bone is directly deposited on a bone surface without previous resorption^10^. This distinction was recently substantiated by demonstrating differences in the chemical composition of the mineral particles, which formed during osteonal remodeling and periosteal modeling, respectively ^11^. Common to all ossification scenarios is that new bone formed by osteoblasts is a non-mineralized collagenous tissue called osteoid ^8, 12^. Besides collagen, osteoid contains water, water binding-proteoglycans, and other non-collagenous proteins (NCPs), such as osteopontin, osteonectin, bone sialoprotein, and osteocalcin ^13, 14^. With a time lag of some days after bone formation, mineral particles start growing in the collagen matrix. This mineralization process starts with a rapid first phase, during which 70% of the final mineral content of the bone is incorporated within a few weeks, followed by a second, slower phase where the mineral content further increases over months and years ^15, 16^. From the investigation of pathological cases, it became clear that the early mineralization phase is critical for creating healthy bone tissue and deviations can lead to hypo-or hypermineralization ^17, 18, 19^. Therefore, investigating the very beginning of physiological mineralization is fundamental to understand how healthy bone is formed and maintained during remodeling. The two processes of remodeling and mineralization running in parallel but lagged and with varying speeds, results in a characteristic spatial-temporal pattern. At a specific time point, the forming osteon consists of a 10-12 μm wide layer of osteoid, followed by a narrow transition zone with a steep gradient in the mineral content, which declines in steepness when approaching the limiting cement line of the osteon. Most importantly, this spatial succession of different mineralization phases allows studying the temporal course of different mineralization stages in a single sample of a forming osteon.

While the function of osteoblasts and osteoclasts in bone remodeling and mineralization is well defined, this holds less when it comes to osteocytes, the most abundant bone cells ^20, 21^. Osteocytes derive from osteoblasts as some of them get encased in the osteoid matrix during bone formation. Late osteoblasts/early osteocytes start to secrete osteocyte-characteristic proteins and form dendritic processes, which eventually pervade the mineralized tissue as well as the osteoid and form a dense cell network. Dendrites and cell body are housed in a pore network of thin (~300 nm in diameter) canals (canaliculi) and ellipsoidal, bulky (~ 300 μm^3^) cavities (osteocyte lacunae) ^22^. A quantification of this lacunocanalicular network (LCN) resulted in staggering values: in human osteons, the average length of canaliculi in a cubic centimeter is 74 km, approximately 80% of the bone is closer than 2.8 μm to the next canaliculus or lacuna, and the total surface of the LCN is about 215 m^2^ in the human body^23, 24^.

Although heavily debated in the past, today there is strong evidence that osteocytes take an active role in Ca and P homeostasis ^25^. Their ability to resorb and secrete material in the neighborhood of the lacunae and canaliculi was observed in lactating mice ^26, 27^, egg-laying hens ^28^, and mechanical unloading^29^, although also lactating mice without traces of osteocytic osteolysis were reported ^30^. The mineral content of the bone was found to be increased in the neighborhood of the LCN ^31, 32^. Osteopontin depositions in the neighborhood of canaliculi might be residuals of the cell’s interaction with its environment ^33, 34^. From a cell-biological point of view, osteocytes are reported to have the ability to secrete proteins, which are known to be involved in the bone formation process ^26, 35^. Furthermore, osteocytes can secrete osteopontin ^36^, matrix extracellular phosphoglycoprotein ^37^, and Dentin matrix protein 1 ^38^, which are known to be inhibitors of mineralization ^39^. Putting together the high density of the LCN and the various interactions between osteocytes and the surrounding extracellular matrix, the question arises if osteocytes are also involved in the early mineralization process, where the first submicrometer-sized electron-dense, mineral agglomerations, the so-called mineralization foci, appear.

The intricate three-dimensional architecture and the sub-micrometer dimensions of the lacunocanalicular network pose a substantial challenge to describe quantitatively how mineralization initiates in spatial relation to the network structure ^40, 41^. The imaging technique of focused ion beam-scanning electron microscopy (FIB-SEM) combines the nanoscopic resolution of electron microscopy with a three-dimensional rendering due to the use of two beams ^42, 43^. Using FIB-SEM, Buss et al. recently investigated sites of bone formation in healthy mice and mice with osteomalacia, a disease resulting in inadequate bone mineralization ^44^. For healthy animals, they report mineralization foci in the sub-micrometer range, which eventually fuse together to form the mineralized matrix. In osteomalacic bone, the foci were much more abundant, over a wider distance creating a less well-defined transition zone. The formation of submicron-sized mineralization foci as being the first mineralization events might be regulated by osteopontin (OPN) as OPN is associated with the foci surface in rat bone ^45^. In the formation of mineralized cartilage, ex-vivo studies on the rat epiphysis suggest that matrix vesicles and proteoglycans are the main mineral nucleation sites ^46, 47^. In bone, although mineralization foci were observed in various studies ^44, 48, 49, 50, 51, 52, 53^, their local distribution with respect to the LCN was never analyzed so far despite one in-vitro study where 20-50 nm wide mineralization foci were associated with the cell processes of MLO-A5 postosteoblast/preosteocyte-like cell line ^54^. In order to investigate the early stage of mineralization in association with osteoblasts/osteocytes, Barragan-Adjemian *et al.* ^54^ conducted an *in vitro* study of MLO-A5, a post-osteoblast/pre-osteocyte cell. They observed that osteoid osteocytes (the immature osteocytes located in the osteoid) and osteoblasts are polarized toward the mineralization front suggesting their activity with respect to the mineralization. Palumbo ^51^ also observed the polarization of osteocytes in the woven bone of chick tibia due to the shifted position of the nucleus toward the vascular side and the higher number of cytoplasmic processes toward the mineralization front. The author suggested that the osteocyte processes toward the mineralization front are involved in the mineralization, and the processes toward vasculature are essential in cell nutrition.

In the present study, we perform a quantitative investigation of mineralization fronts at forming osteons of healthy women. Our aim is to localize the very first mineralization events and to describe their temporal development in relation to the cell network present in osteoid. We study nine different forming osteons using contrast-enhanced FIB-SEM imaging. Iodine staining of the bone samples, to specifically increase the electron density of the organic matrix (and thus the backscattered electron yield), allows us to discriminate osteoid, mineralized tissue, and the LCN pervading both tissue types. A spatial resolution of 40 nm in three-dimensional FIB-SEM imaging provides access to the early mineralization event of foci formation.

## Results

### Three-dimensional heterogeneity of mineral content in a forming osteon

Figure 1a shows the result of a typical FIB-SEM measurement. Extracted from the three-dimensional image stack, the two backscattered electron (BSE) images from two planes perpendicular to each other represent the FIB-SEM dataset where the brightness of voxels can be related to the local mineral content (also see Supplementary Movie 1). The imaged volume shows a forming osteon with unmineralized matrix (dark voxels) in the vicinity of the Haversian canal, and a higher mineral content (bright voxels) further away from the canal. At the transition between unmineralized and mineralized tissue, whitish speckles – the mineralization foci – can be observed. Based on the gray value probability distribution (GVPD) of all voxels, a distinction was made between the unmineralized and mineralized matrix (Fig. 2, see details about the image segmentation in the Methods section). The osteocyte lacuna is located in the mineralized tissue, and numerous canaliculi connect the lacuna to the surface of the Haversian canal. Figure 1b shows an image of the FIB-SEM dataset taken from a transversal cross-section through the osteon. Reading the figure from left to right, the Haversian canal (brown), the unmineralized (gray) and mineralized tissue (whitish) with the transition zone in-between are visible as expected from a forming osteon ^54^. Figure 1c is a zoom into the transition zone, where the image plane is now perpendicular to the plane of Figure 1b and roughly tangential to the transition zone. The green dots represent cross-sections through the canaliculi with a diameter of roughly 0.5 μm. Thus the 3D image sets comprise two types of canals of very different dimensions, the Haversian canal at the micro-scale and the canaliculi at the nano-scale. In the following, we analyze the mineral content of the tissue as a function of the distance to these canals (in Figure 1b and 1c, the direction of the increase of distance from the surface of the canals is shown by red arrows).

**Fig. 1:**
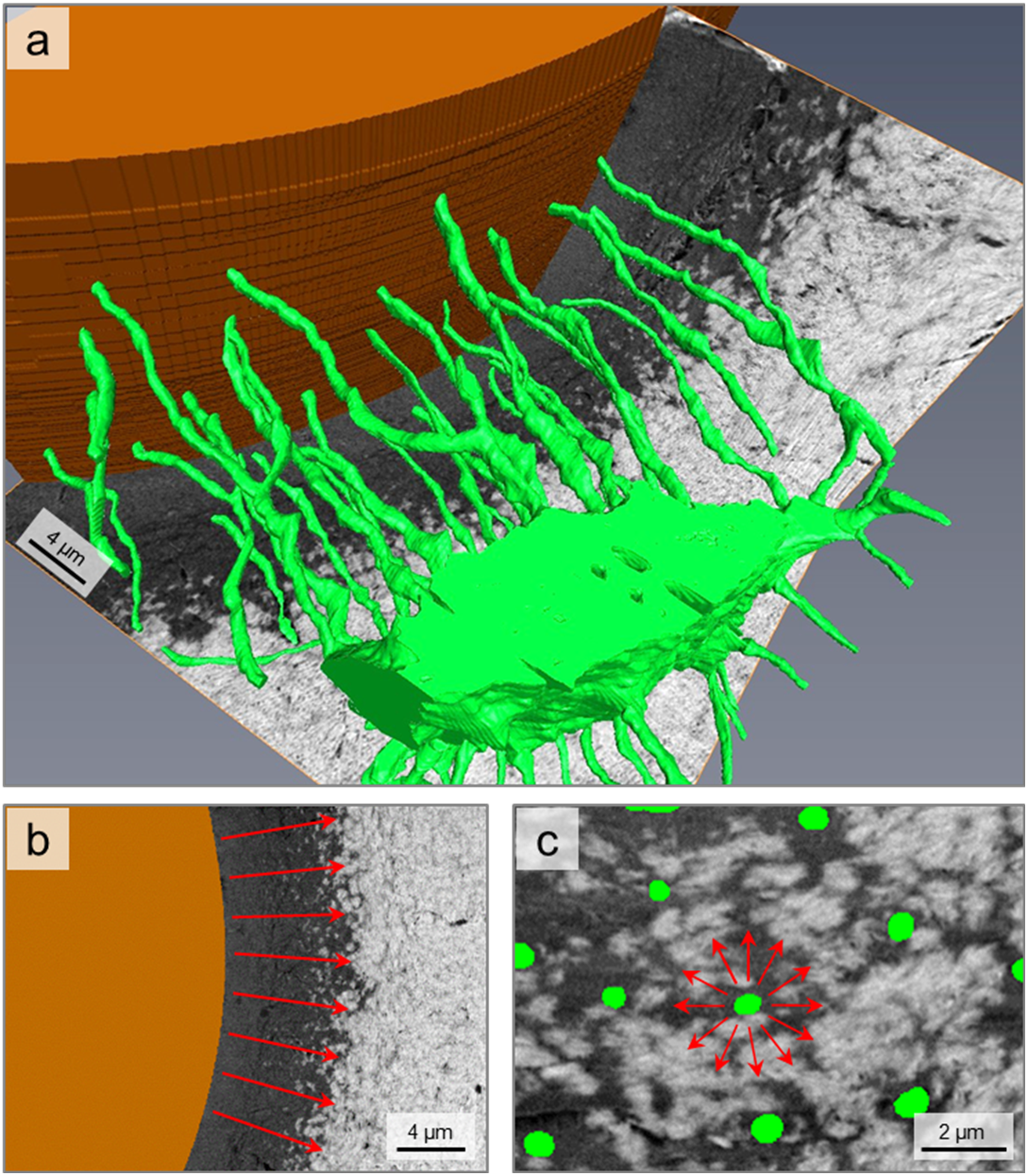
Overview of the osteocyte network within an osteon. a) 3D rendering of an osteocyte lacuna, emerging canaliculi (green), and the Haversian canal (brown) within a forming osteon. The perpendicular cutting planes are FIB/SEM backscattered electron images showing mineralized matrix (bright voxels) and unmineralized matrix (dark voxels). b) Cross-section of the Haversian canal with osteoid and the transition to mineralized tissue. Within the transition zone, mineralization foci are visible as bright “islands” in a dark unmineralized matrix tissue. c) Cross-section perpendicular to the direction of canaliculi so that canaliculi appear as round dots (green). The red arrows in panel b and c show the direction of increase in distance from the Haversian canal and canaliculus surface, respectively.

**Fig. 2:**
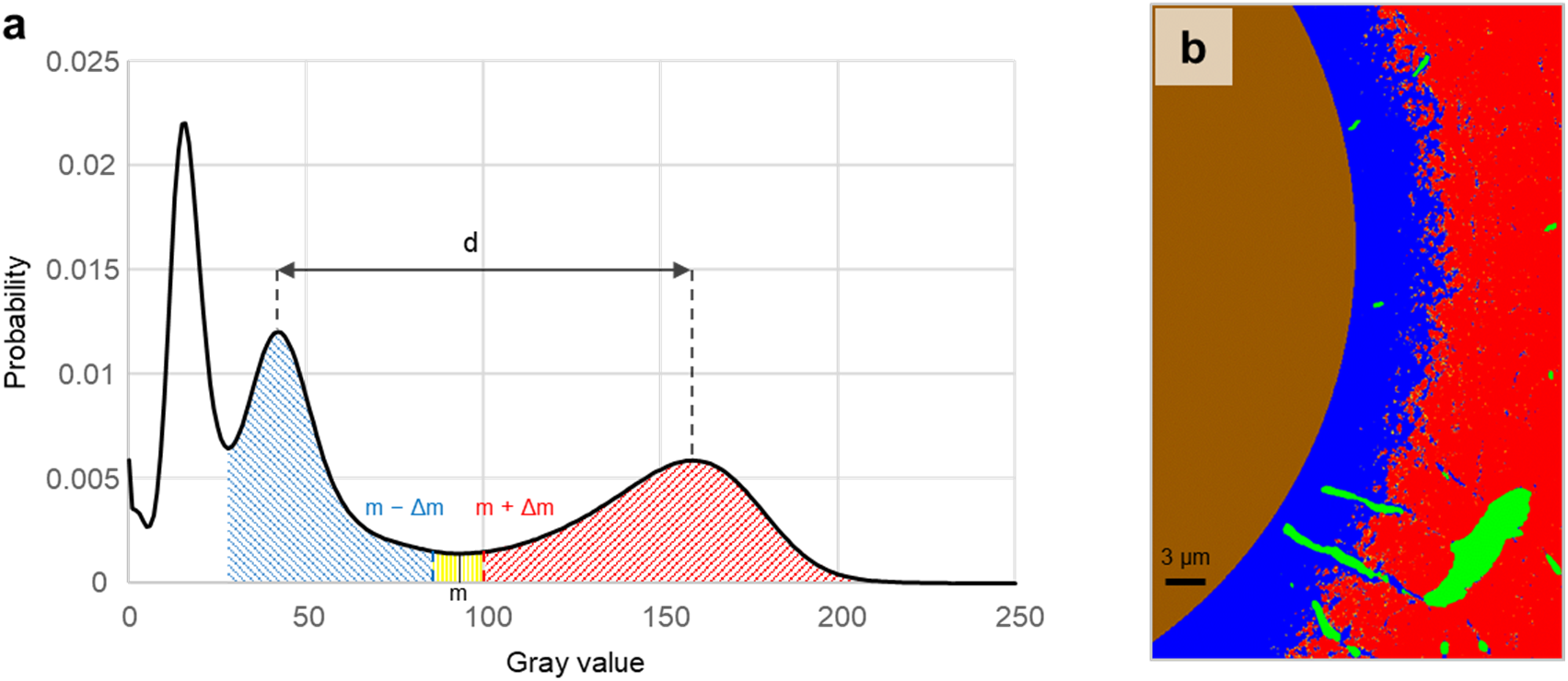
Thresholding and segmentation of the bone tissue. a) Segmentation of unmineralized (blue area) and mineralized (red area) matrix based on the gray value probability distribution (GVPD) of the 3D data sets. The minimum between the two peaks at higher gray values is denoted by m, the distance between the maxima of the peaks by d. Voxels with gray values above m + Δm and below m – Δm, where Δm = 0.05*d, are considered to belong to the mineralized or unmineralized matrix, respectively. The sharp peak at the lowest gray values corresponds to PMMA and thus is linked to regions previously assigned in the images to the PMMA phase via manual segmentation (see details in the Methods section). b) Representative slice of the 3D dataset as result of the segmentation. Four different compartments are distinguished: (i) Haversian canal (brown); (ii) osteocyte lacunae and canaliculi (green); (iii) non-mineralized matrix (blue); and (iv) mineralized matrix (red). For the analysis of mineralization, the volume fraction of mineralized matrix per total matrix (V_MM_/V_TM_) was calculated based on the dividing of the red area by the red, yellow, and blue area.

### Mineral content as a function of distance from the Haversian canal

After the segmentation of the Haversian canal, a distance transform was used to assign each voxel in the image the shortest distance from the Haversian canal. For all voxels with the same distance from the Haversian canal (used bin size was 0.27 μm), a gray level frequency distribution was calculated. Normalization by the total number of voxels at a given distance results in a gray value probability distribution (GVPD) (see Materials and Methods for details). Consequently, the color code in Figure 3a reveals the likelihood to find, at a given distance from the Haversian canal, a voxel with a specific gray value. For the analyzed dataset shown in Figure 1, the clear majority of voxels have low gray values for distances smaller than 6 μm from the Haversian canal. For a distance larger than 9 μm this majority changes to a much larger gray value. Considering the variability of the gray values for a specific distance, the variability is smaller for short distances from the Haversian canal as indicated by the dark narrow “band” (Fig. 3a). Quantification of the transition between the low and high gray value was obtained by calculating the average gray value for each distance from the Haversian canal, which resulted in the black line of Figure 3a.

**Fig. 3:**
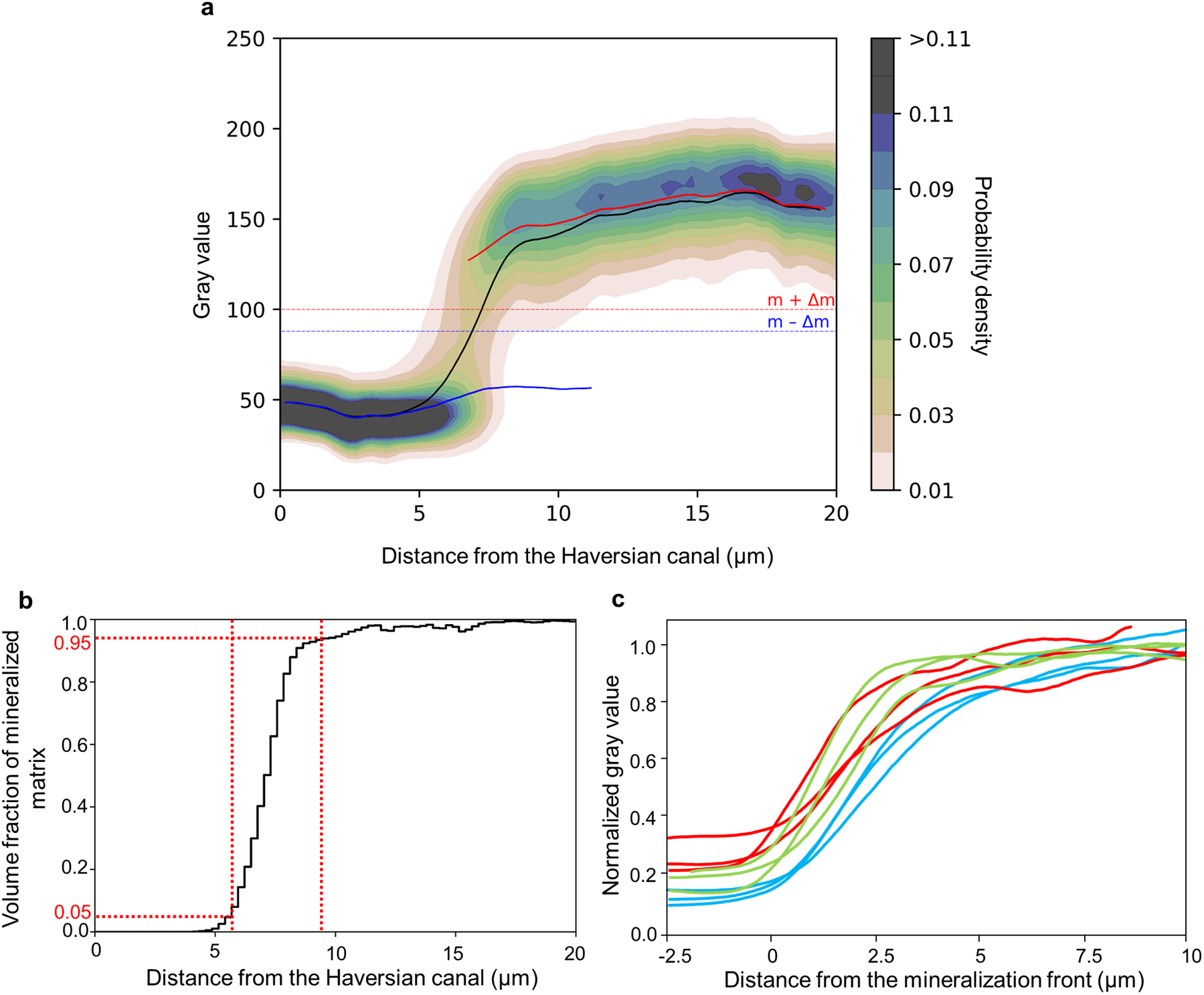
Bone mineralization with respect to the distance from the Haversian canal. a) Gray value probability distribution (GVPD) as a function of the distance from the Haversian canal (osteon #2, sample #1). For the graph, a 3D subvolume (~ 20×25×40 μm^3^) of the FIB/SEM dataset was analyzed. The color code provides the likelihood to find a voxel with a given gray value at a specific distance from the Haversian canal (i.e., the sum of all probabilities for each distance bin sums up to 1, bin width was equal to 0.27 μm). The gray values m-Δm and m+Δm define the thresholds for unmineralized and mineralized matrix, respectively (see also Fig. 2a). The smooth curves are the moving averages of the gray values (using a window width of 5% of all data points) for the whole matrix (black), the unmineralized (blue), and mineralized matrix (red), respectively. b) The volume fraction of mineralized matrix per total matrix (V_MM_/V_TM_), i.e. the fraction of mineralized voxels per total voxels at each distance bin, plotted as a function of the distance from the Haversian canal for the same osteon as in Figure 3a. The vertical red dashed lines mark the start and end positions of the transition zone defined at positions where 5% and 95% of the matrix is mineralized. The start point of the transition zone is referred to as the mineralization front and is used as a positional reference. c) Normalized average gray values of the nine osteons as a function of the distance to the mineralization front. For the normalization, the peak positions in the gray level distribution (Fig. 2a) corresponding to PMMA and to the mineralized matrix were set equal to 0 and 1, respectively, by the following linear equation: 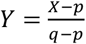, where Y is the normalized gray value, X is the original gray value, p is the gray value of maximum frequency of PMMA, and q is the gray value of maximum frequency of mineralized matrix. The green, blue, and red curves correspond to the osteons of samples #1, #2, and #3, respectively.

The average gray value displays a sigmoidal shape with a steep transition between two roughly constant plateau values. For high gray values, a trend for a further increase of the average gray value when moving away from the Haversian canal can be observed. However, the slope of the curve is substantially smaller compared to the transition between low and high gray values.

We evaluated the average value for unmineralized and mineralized matrix separately, where we deliberately excluded voxels with an “intermediate” gray value (i.e., between *m* — *Δm* and *m* + *Δm* in Fig. 2a). The blue and the red curve in Fig. 3a represent the average value for the unmineralized and the mineralized matrix, respectively. The horizontality of the blue curve excludes a movement of the peak. However, the red curve representing the mineralized matrix displays a gentle slope indicating that the peak is moving to larger gray values while building up.

In Figure 3b the volume fraction of mineralized matrix per total matrix (V_MM_/V_TM_) was plotted as a function of the distance from the Haversian canal for the same osteon as in Fig. 3a. This way one can precisely quantify the extension of the transition zone in three dimensions. The start and end points of the transition zone were defined as the positions where 5% and 95% of the matrix is mineralized, respectively (vertical red dashed lines in Fig. 3b). The start point at 5% was also used in the following as defining the reference position of the mineralization front. From the evaluation of all nine osteons, the width of the transition zone was determined to be 3.67 ± 0.57 μm.

The definition of the position of the mineralization front allowed to align the average gray value as a function of the distance from the Haversian canal for all nine osteons in a single plot (Fig. 3c). For this plot, the gray values were normalized in a way that the peak position of the PMMA phase was set equal to 0 and the peak position of the mineralized matrix was set equal to 1. The average gray value shows a sigmoidal shape for all osteons.

### Mineral content as a function of distance from canaliculi

Using the same method of analysis as for the mineralization relative to the Haversian canal, but now on a much smaller length scale, the mineralized and unmineralized matrix around canaliculi was quantitatively described. To take into account that the tissue in a forming osteon has a very heterogeneous mineral content, a cuboid region parallel to the mineralization front with a quadratic cross-section of the size of 8×8 μm^2^ was selected and divided into three subvolumes depending on the volume fraction of mineralized matrix per total matrix (V_MM_/V_TM_, see Fig. 2a): lowly mineralized tissue with a V_MM_/V_TM_ between 5% and 35%, medium mineralized tissue with a V_MM_/V_TM_ between 35% and 65%, and highly mineralized tissue with a V_MM_/V_TM_ between 65% and 95%. Figure 4a-c show three two-dimensional BSE images taken from the lowly, medium, and highly mineralized tissue. It can be observed that the green round dots marking the cross-sections of canaliculi are situated in regions of low mineral content. This is true even for the highly mineralized tissue (Fig. 4c), where most of the tissue is mineralized except the close environment of the canaliculi.

**Fig. 4:**
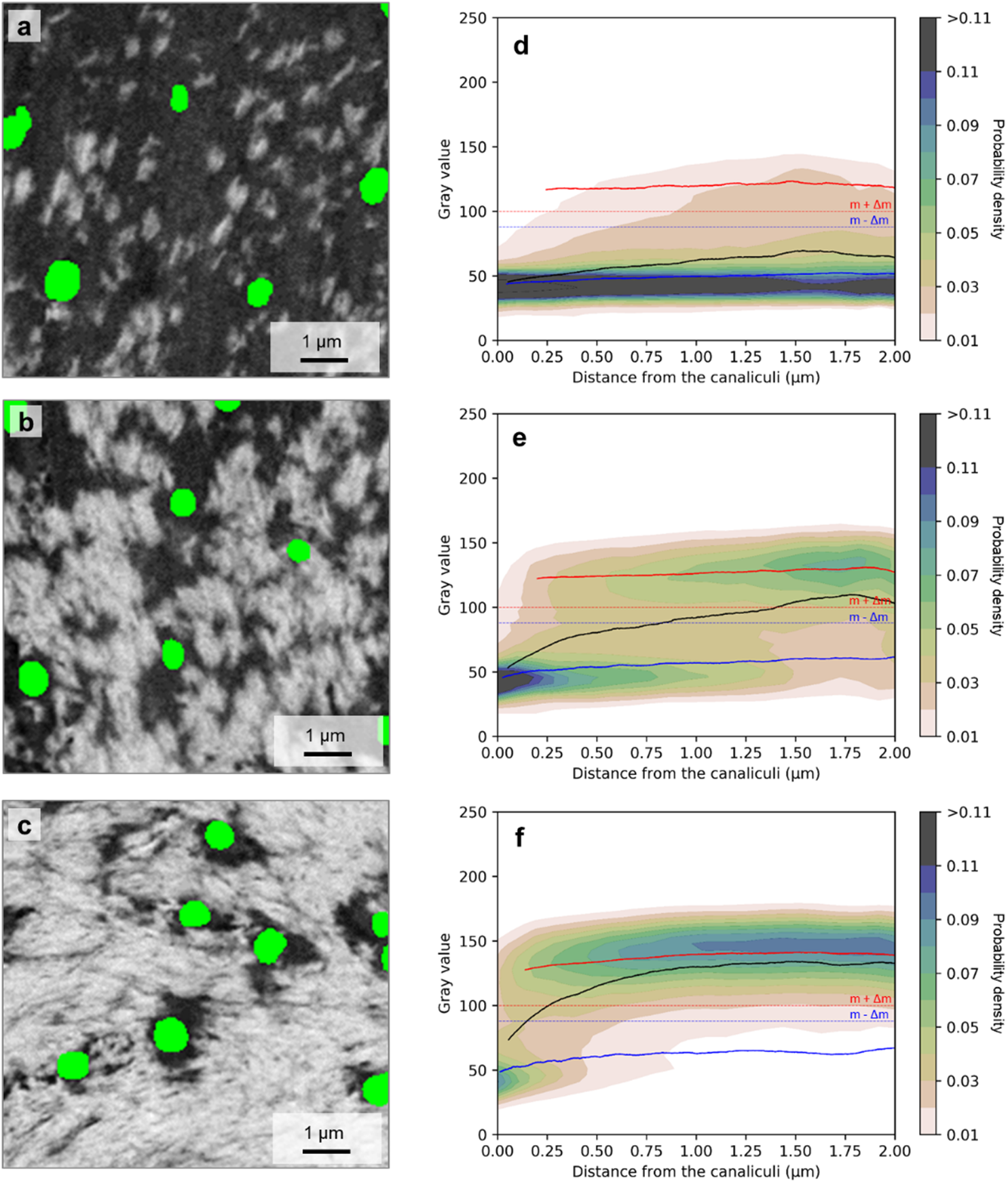
Bone mineralization with respect to the distance from the canaliculi. Backscattered electron (BSE) images of the unmineralized (dark) and mineralized matrix (bright) around canaliculi (green dots) in a) lowly mineralized (V_MM_/V_TM_: 5% - 35%), b) medium mineralized (V_MM_/V_TM_: 35% - 65%), and c) highly mineralized tissue (V_MM_/V_TM_: 65% - 95%). The image plane is roughly perpendicular to the direction of canaliculi, which causes their almost circular appearance. Gray value probability distributions are shown as a function of the distance from the canaliculi for d) lowly, e) medium and f) highly mineralized tissue. The black line corresponds to the overall average gray value, while the blue and red lines correspond to the average value of the unmineralized and mineralized tissue, respectively (see Fig. 2a for the definition of unmineralized and mineralized tissue). Note that the probability distribution shows the evaluation for the subvolumes of the lowly, medium, and highly mineralized tissue of an osteon (osteon #2) of sample #1 and not for the single BSE images shown to the left.

As before for the whole osteon, Figure 4d-f show the gray value probability distributions (GVPDs), this time as a function of the distance from the canaliculi. The black line corresponds to the average gray value, while the blue and the red line are the average gray value when the evaluation is restricted to the unmineralized and mineralized matrix, respectively. For all three tissues (lowly, medium, and highly mineralized) the lowest gray values are found close to the canaliculi, and the average gray value increases when moving away from the canaliculi. For the highly mineralized tissue (Fig. 4f) again a transition occurs from an unmineralized matrix around the canaliculi to a mineralized matrix further away from the canaliculi. The constant values over the distance of the blue and the red curves demonstrate that the transition occurs by a change in the volume fraction between mineralized and unmineralized matrix and not by a gradual shift of the gray values in the unmineralized matrix. The reported results refer to an osteon from sample #1 but similar patterns were also observed in all other datasets as shown in Supplementary Figure 2.

### Estimated time-evolution of the transition zone on the canalicular scale

In an analysis, which included data from all nine investigated osteons, our aim was to quantify the transition zone from mineralized to unmineralized matrix around canaliculi. This zone was characterized within subvolumes defined by their volume fraction of mineralized matrix (V_MM_) per volume of total matrix (V_TM_). We use V_MM_/V_TM_ to define stages of tissue mineralization and in addition to the three different stages defined above (lowly, medium, and highly mineralized), we included the extreme cases of unmineralized (V_MM_/V_TM_ = 0%) and fully mineralized tissue (V_MM_/V_TM_ > 99.9%) as reference points, plus an almost fully mineralized tissue (with V_MM_/V_TM_ > 95%). Approximate values of the distance from the starting mineralization front where these 6 successive stages are found are indicated in the left panel of Figure 5. The average volume fraction of the mineralized matrix was evaluated in ring-like volumes (ring width = 0.095 μm) with a specific distance to the nearest canaliculus (D = 0.1 μm, 0.25 μm, 0.5 μm, 2 μm). Moving away from the Haversian canal with a continuous increase in the mineral content, the mineralization process far away from the canaliculus (D = 2 μm) sets in first and reaches for highly mineralized tissue (stage III) already a value of 90 % for the V_MM_/V_TM_ (dark brown data point). At the same stage III, a region close to the canaliculus (D = 0.1 μm) displays a V_MM_/V_TM_ of only 20% (bright flesh-colored data point). For a region that close to the canaliculus, mineralization sets in at later stages and reaches values above 90% V_MM_/V_TM_ only in the limiting case of fully mineralized tissue.

**Fig. 5:**
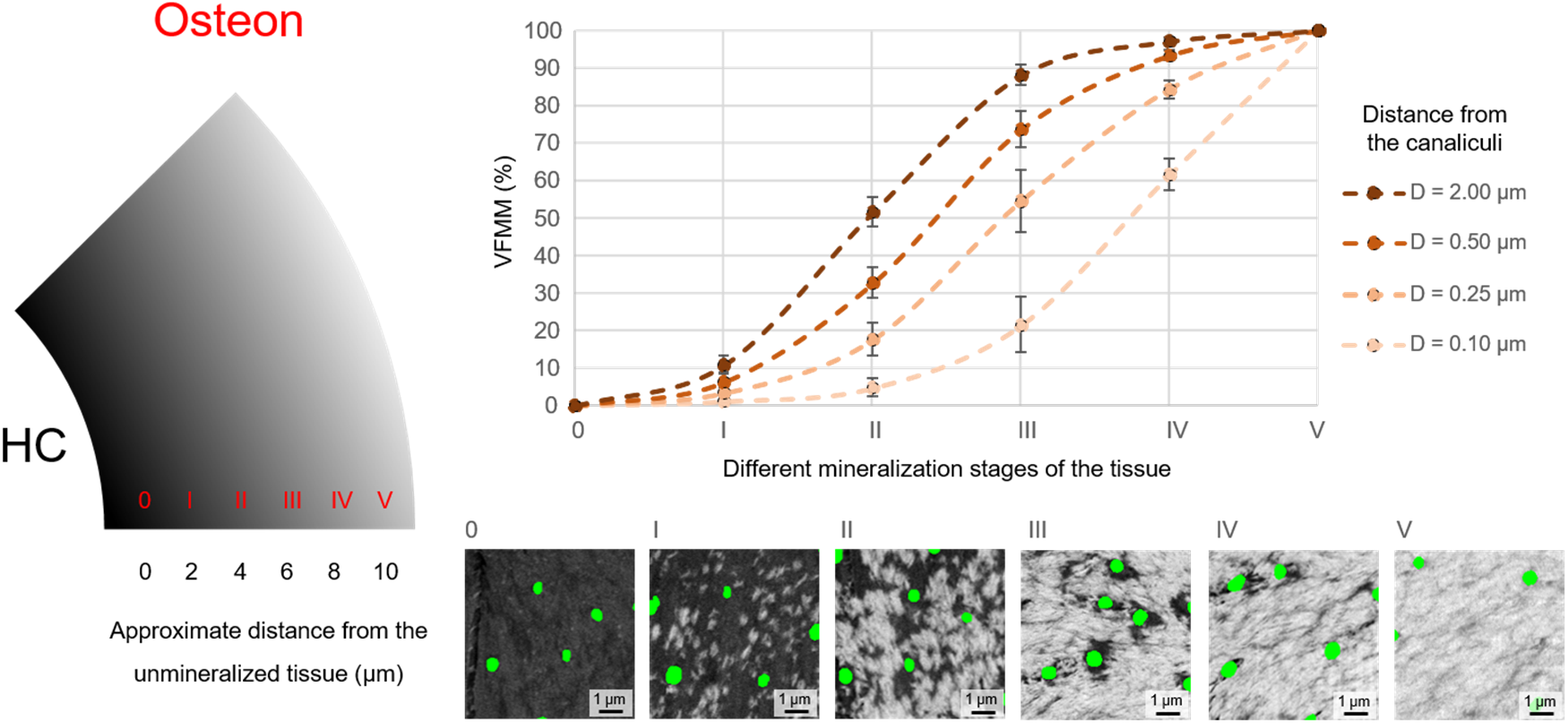
Estimated time-evolution of the transition zone around the canaliculi front. Based on its overall volume fraction of mineralized matrix per total matrix (V_MM_/V_TM_), the tissue within an osteon (see left of figure, HC being the Haversian canal) was subdivided into six stages (0: unmineralized, V_MM_/V_TM_ = 0%, I: lowly mineralized (V_MM_/V_TM_: 5% - 35%), II: medium mineralized (V_MM_/V_TM_: 35% - 65%), III: highly mineralized (V_MM_/V_TM_: 65% - 95%), IV: almost fully mineralized (V_MM_/V_TM_ > 95%), V: fully mineralized (V_MM_/V_TM_ > 99.9%, at far distance from the mineralization front), which correspond to temporal stages of progressing mineralization. Data points show the average V_MM_/V_TM_ at different distances from the canaliculi (bright flesh-colored = 0.1 μm, flesh-colored = 0.25 μm, brown = 0.5 μm and dark brown = 2 μm; evaluation in a ring-like volume of ring width of 0.095 μm). Values were obtained by averaging over all nine investigated osteons and error bars denote the standard error.

### The size distribution of the mineralization foci

In the early phases of the mineralization process, the FIB/SEM images allow a distinction of well-separated mineralization foci. In lowly mineralized tissue, we quantified the formation and growth of these mineralization foci by analyzing their sizes. For each focus, the local volume fraction of mineralized matrix per total matrix (V_MM_/V_TM_) was calculated in a cube of 1 μm^3^ with the focus at its center. Depending on the V_MM_/V_TM_ of their surrounding matrix, the foci were then allocated into one of three groups: V_MM_/V_TM_ of 0.1 – 5% (Fig. 6b), V_MM_/V_TM_ of 5 – 10% (Fig. 6c), and V_MM_/V_TM_ of 10 – 20% (Fig. 6d). For these three groups, Figure 6b-d show the contribution to the mineral volume fraction of different-sized foci as a function of the distance to the closest canaliculus, i.e. the color code denotes the probability weighted by the foci volume to find a focus with a specific volume at a specific distance from the canaliculi.

**Fig. 6:**
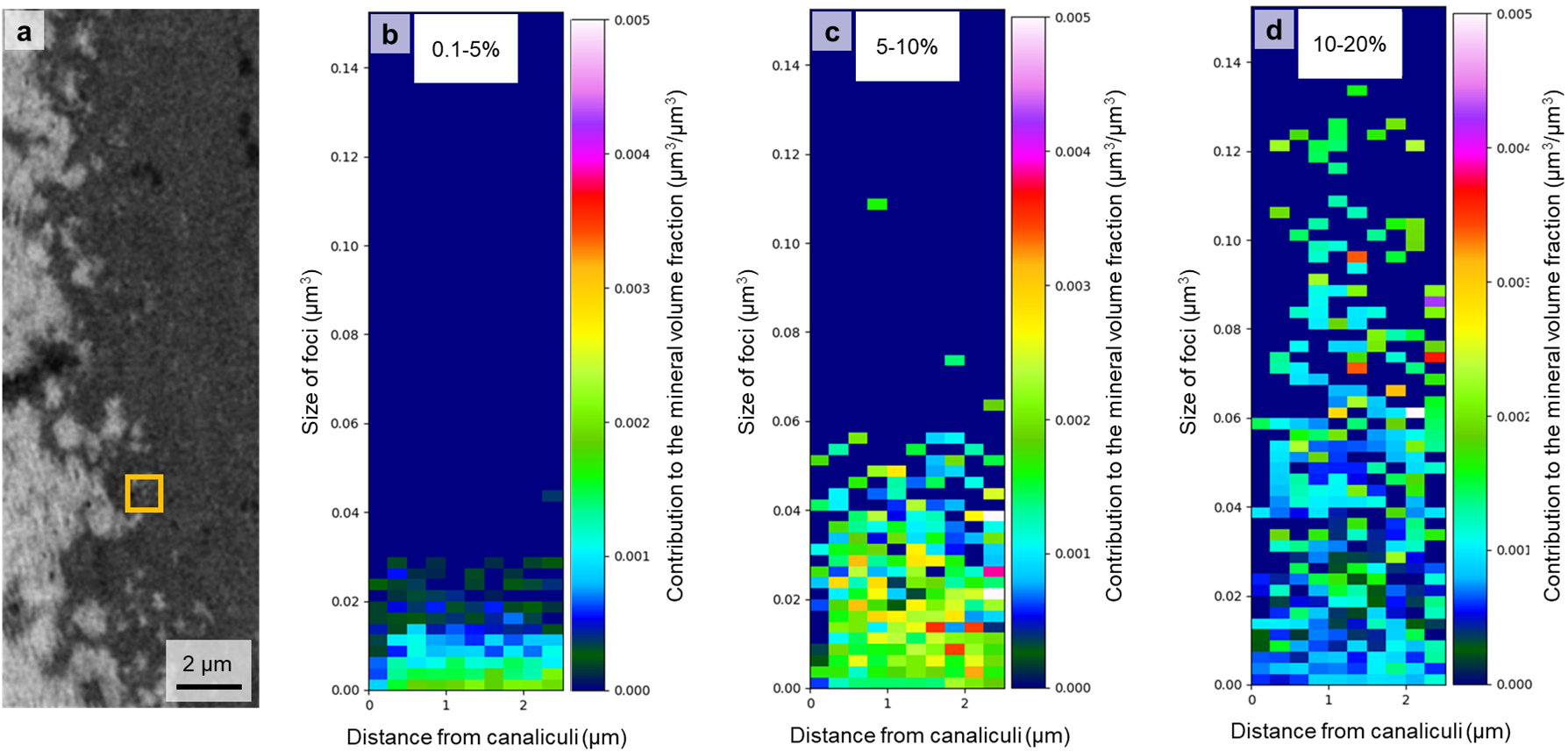
Probability distribution of mineralization foci volume as a function of the distance from the canaliculi. a) The local volume fraction of mineralized matrix per total matrix (V_MM_/V_TM_) was calculated within a 1 μm^3^ cube surrounding the foci (illustrated by a yellow square). Depending on the V_MM_/V_TM_ of the surrounding matrix, foci were allocated into one of three groups: V_MM_/V_TM_ = 0.1% - 5%, 5% - 10%) and 10% to 20%. b-d) The color in the 2D-histograms denotes how different-sized foci contribute to the mineral volume fraction for a given distance to the closest canaliculus, i.e. the probability weighted by the foci volume to find a focus with a specific volume at a specific distance from the canaliculi. Values were obtained over all nine investigated osteons.

Figure 6b shows that formation of mineralization foci starts more or less independent of the distance from the canaliculi (Fig. 6b), except for the zone close to the canaliculi (distance < 250 nm), where the formation probability of foci is low. Comparison with the situation at a higher mineral volume fraction (Fig. 6c) reveals a growth of the foci in volume, but their sizes remain roughly independent of the distance from the canaliculi. The only observed trend is that the number of growing foci is larger farther away from the canaliculi. With the progression of mineralization (Fig. 6d) the formation of new small foci is decreasing, while the existing ones are continuing to grow. Coalescence of foci results in instantaneous volume increases to values, which are beyond the scale of the y-axis chosen in Figure 6d.

## Discussion

We used FIB-SEM to image the course of mineralization in a newly forming osteon with nanometer resolution and in three dimensions. We observed two mineral gradients at two different levels, the micrometer scale around the Haversian canal (diameter ~50 μm) and the submicrometer scale more than two orders of magnitude smaller, around the canaliculus (diameter ~0.3 μm).

The transition zone from unmineralized to mineralized bone in a forming osteon was previously described using electron microscopy, X-ray scattering, and spectroscopic methods in 2D ^11^. Our 3D imaging method allows us to quantify the width of the transition zone to be 3.67 ± 0.57 μm. A similar transition zone has been observed by Buss et al.^44^ in healthy mouse bone with a much narrower width. The roughly 5 μm thick osteoid seam surrounding the Haversian canal is suggested to be kept free of mineral by circulating mineralization inhibitors, such as pyrophosphate, or local inhibitors such as osteopontin, that suppress mineralization close to the osteoblasts ^53, 55^. To initiate mineral formation within the transition zone, enzymes such as TNAP and PHEX were proposed to remove the effect of inhibitors ^55^. Since the body fluid is rich of Ca and PO4 ions that tend to crystalize ^56^, these inhibition and promotion mechanisms are particularly important to trigger the mineralization where it is desired and to avoid ectopic mineralization. During progression of mineralization, the transition zone is moving towards the Haversian canal. Far away from the canal is the location where the first fully mineralized tissue can be found.

The high resolution of our images showed that the transition between unmineralized and mineralized bone is not continuous with a homogeneous mineral content at a given distance from the Haversian canal. Instead, a closer look revealed a granular nature of the transition zone with discrete mineralization foci. This already previously observed granularity ^54^ can now be quantitatively described based on 3D imaging.

To understand the role of the osteocyte network in early mineralization, we evaluated size and location of the first mineralization foci with respect to the lacunocanalicular network. The spatial correlation of the foci with the LCN structure indicates some involvement of osteocytes in mineralization. Most importantly, this spatial correlation is not such that the local mineral content decreases from the canaliculi, but it rather increases. Hence, the matrix surrounding canaliculi is comparatively poor of mineral in matrix undergoing mineralization. We refer to this zone of low mineral content around canaliculi as “halo zone” (Fig. 4). Our image quantification demonstrated a strongly reduced probability of foci formation in the halo zone compared to regions further away from the canaliculus (Fig. 6). This observation excludes simple diffusion of mineral precursors originating from the canaliculi, since then one would expect a Gaussian decrease of concentration with distance from LCN.

As suggested for the mineral-free osteoid around the Haversian canal, also the halo zone around canaliculi might be explained by the action of mineralization inhibitors. It is known that inhibitors of mineralization are secreted by osteocytes, which suppress mineralization too close to the cell surface. We do not have evidence for the specific molecule(s) involved in this process, but we interpret our experimental finding as the result of a co-diffusion of promotors (i.e., mineral precursors) and inhibitors of mineralization from the LCN. Such an interplay between diffusants is reminiscent of the Turing model ^57^– and more generally described in the mathematical framework of reaction-diffusion systems–which are known to generate complex patterns in various biological (and chemical) systems, depending on geometry and the relative diffusivity of the diffusants. In addition to these mobile players, it is not unlikely that spatially fixed molecules influence the course of early mineralization. Collagen was proposed to act as a nucleator^58^, while macromolecules like proteoglycans could act as practically immobile inhibitors. While it is beyond the scope of the current work to develop a quantitative version of a Turing-like model with so many interacting players, we are confident that the experimentally observed patterns can be obtained with a suitable set of model parameters, guided by detailed analyses of such models that can be found in literature ^59, 60^.

The sketch of Figure 7 summarizes how we imagine the mineralization process in a forming osteon in relation to the network of canaliculi. For reasons of simplicity, in the longitudinal section of Figure 7a, only a single canaliculus is depicted, which bridges the tissue between the osteocyte body and the Haversian canal. The osteocyte body and part of the canaliculus are within the mineralized tissue beyond the unmineralized osteoid. The canaliculus, or more likely the osteocyte cell process within the canaliculus, is the source of mineralization precursors and inhibitors (shown as curled purple structures), which diffuses into the matrix building up the usual diffusion gradient with a high concentration close to the source and rapid decrease in case that the inhibitor diffuses slowly. For this reason, the actual precipitation of mineral is likely to occur further away from the canaliculus where the inhibitor concentration is lower. With more mineralization precursor arriving in the matrix, its concentration will eventually outperform the inhibitors even closer to the canaliculus, so that the mineral front approaches the canaliculus with increasing overall mineral content. While at some point the formation of new foci ceases, the growth of already existing foci continues until they reach the surface of the canaliculus and thereby filling in the halo zone. This closing in of the mineralized region towards the canaliculus is illustrated in the transversal cross-sections of Figure 7b.

**Fig 7:**
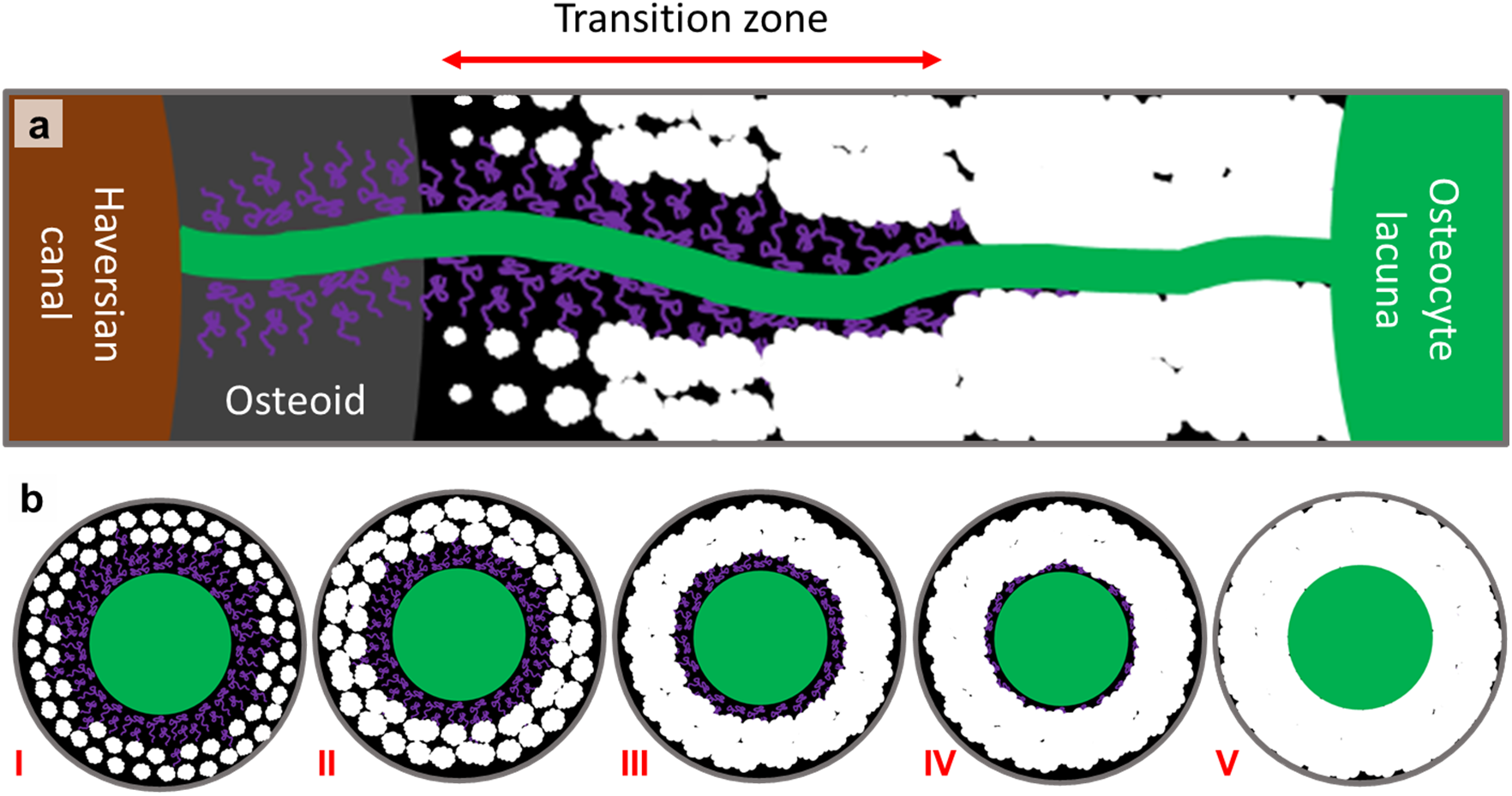
Mineralization pattern around the canals in a forming osteon. a) Sketch of the course of mineralization with respect to the Haversian canal and the canaliculi. The green canal is a canaliculus and the bright spots are the mineralization foci. The curled purple structures represent mineralization inhibitors. The newly formed foci are small, but later on, they become bigger and fuse together to create a fully mineralized matrix. Note that the Haversian mineralization front and the mineralization foci are depicted larger than their real size for better visualization. b) Transversal section through a canaliculus showing the distribution of foci around the canaliculus at different stages with i) low, ii) medium and iii) high volume fraction of mineralized matrix per total matrix (V_MM_/V_TM_). At stage iv, the halo zone is almost filled in and at stage v, V_MM_/V_TM_ is almost 100%. The inhibitors are not to scale.

The current work reveals a lowly mineralized region around canaliculi in the transition zone between osteoid and mineralized matrix indicating that the osteocytes play a significant role in the early mineralization process. In contrast to that, studies on anosteocytic fish bones show that physiological mineralization happens in the absence of osteocytes and thus without LCN ^61^. On the other hand, the even narrower porosity network (with a diameter of ~40 nm) discovered in turkey leg tendon suggests strong cellular (in this case tenocytes) control of the mineralization process ^62^. Hence, the question rises how universal or diverse mineralization mechanisms are between species or even between sites within the same animal ^63^. We see a high demand for comparative studies to reveal similarities and differences in the mineralization processes in order to elucidate the fundamental biochemical mineralization mechanisms and how they are governed by cell activity. We suggest that one of these basic mechanisms is the co-diffusion of mineral precursors, promotors, and inhibitors. To characterize such a system, Turing models describing reaction-diffusion processes can potentially be investigated as they define a framework to describe pattern formation involving various re-or counter-acting diffusants.

## Methods

### Sample preparation

The presented data were obtained from three femoral midshaft necropsy samples of middle-aged women (48 (sample #1), 55 (sample #2), and 56 (sample #3) years old) without any known bone-related disease. All samples were provided by the Department of Forensic Medicine and the Department of Anatomy of the Medical University of Vienna according to ethic commission regulations of the university (EK no. 1757/2013). For storage, the native bones were frozen at −20 °C. Sectors were cut out of the cortical ring from midshaft samples and thawed prior to dehydration with an ethanol series. Subsequently, the specimens were put into a falcon tube filled with a 40 ml rhodamine6G/ethanol solution (0.417g/100ml) for 72 hours while renewing the staining fluid twice following an established protocol^31^. The stained samples were then embedded in polymethylmethacrylate (PMMA) as described in ^64^. During this procedure, PMMA fills not only the large porosity of the Haversian canals, but also the much smaller LCN porosity. Using a low-speed diamond saw (Buehler Isoment, Lake Bluff, Illinois) the sample blocks were cut transversally (i.e., perpendicular to the long axis of the bone) to obtain a slice of 1 mm thickness with osteons exposed at the surface with a roughly circular shape. One of the sample surfaces (~0.5 x 0.5 mm^2^) was then ground with sandpaper of descending grain size and polished with 3 and 1 μm grain size diamond suspension (Logitech PM5, Glasgow, Scotland).

### Iodine staining of the unmineralized tissue

To be able to discriminate between unmineralized matrix and embedding material using backscattered electron microscopy, contrast-enhancing methods are needed as otherwise, the average atomic numbers of the two materials are too similar (close to 6, i.e. the atomic number of carbon). Since our research strategy is to use FIB-SEM for the acquisition of 3D datasets with distinguishable canaliculi in the mineralized and the unmineralized matrix, a stain is needed that penetrates the sample homogenously at least up to a typical field of view depth of 25 μm. Boyde et al. described a procedure for enhancing the SEM backscatter contrast in soft tissue in 2D by exposing a PMMA embedded bone sample block to iodine vapor ^65, 66^. Based on this finding, we developed an iodine vapor staining protocol that provides appropriate staining results: 0.1 mg solid iodine (Alfa Aesar, Karlsruhe, Germany) were placed in a 25 ml autoclave sealed glass container (Duran, Wertheim/Main, Germany) next to the PMMA embedded polished sample for six days followed by a second identical staining cycle. We observed that the embedding material remained unstained and that only in the unmineralized osteoid the electron density (and, therefore, the backscattered electron coefficient) increased due to the incorporation of iodine with its high atomic number of 53.

### Focused ion beam-scanning electron microscopy (FIB-SEM)

ROIs located at forming osteons (defined using pre-characterization by ESEM and CLSM, see Supplementary Figure 1) were imaged with FIB-SEM (Crossbeam 540 Zeiss, Oberkochen, Germany) obtaining 3D datasets with typical dimensions of ~40×30×50 μm^3^ with an isotropic voxel size of ~40 nm. Prior to FIB-SEM imaging, the samples were carbon coated with a vacuum carbon evaporator sputter coater (CED 030, Bal-tec/Balzers, Liechtenstein, Germany) to reduce charging effects. To avoid sample shrinkage during the measurement, the samples were mounted inside the vacuum chamber at least 24 hours before imaging. The working distance during the serial surface imaging was set to the coincidence point of the FIB and the SEM beams, i.e. at around 5.1 mm. The milling process was conducted with high- and low-power gallium ion probes (65 nA and 0.7 nA at 30 kV acceleration voltage, respectively). With the higher power, a large hole (~100×100×100 μm^3^) was milled, while the latter was used during the serial surface imaging process featuring a slice thickness of 40 nm and a milling depth of 50 μm due to ~5 seconds milling time per slice. High-resolution secondary and backscattered electron images (BSE) (~40×40 nm^2^) were obtained with 2.5 kV acceleration voltage, 1 nA beam current, and 28 s acquisition time per frame using 40 averaging cycles per line. Brightness and contrast were set to avoid over-or underexposure of regions in the backscattered electron images. Post-processing of the BSE image stacks was performed using in-house developed Python scripts to take into account (i) misalignment/sample drift, (ii) image noise, and (iii) curtaining artifacts originating from the milling process.

### Image segmentation

The 3D FIB-SEM datasets were segmented into different compartments based on structural features and backscattered electron image brightness in a two-step process using Amira (Version 2019.38, Zuse Institute Berlin (ZIB), Berlin, Germany). First, manual segmentation was performed to distinguish between PMMA (corresponding to Haversian canal, osteocyte lacunae, and canaliculi) and bone. Since PMMA is not stained by iodine and, therefore, the darkest compartment in the datasets, the Haversian canals could easily be identified due to the clear contrast to the adjacent unmineralized iodine stained osteoid tissue. Osteocyte lacunae and canaliculi in the mineralized and unmineralized tissues were manually traced every 5 slices according to their typical shape and dark appearance followed by an interpolation of the remaining slices.

In a second step, the bone region was further sub-divided into regions of mineralized and unmineralized matrix based on the analysis of the dataset’s probability distribution of the gray values in the dataset (see Fig. 2a). To consider for variations in the iodine staining intensity and for differences in the brightness and contrast settings during image acquisition, for each dataset the minimum of the probability distribution (m) between the peak of the mineralized and the unmineralized matrix and the gray value difference (d) between the two peak positions was determined. As gray values close to m (Fig. 2a, yellow) could not clearly be assigned to one of the two phases due to noise or partial volume effects, only gray values above m + Δm and below m – Δm are considered to belong to the mineralized (red) or unmineralized (blue) matrix, respectively. Δm was chosen to be 5% of d. A representative 2D slice of a segmented 3D volume is shown in Fig. 2b.

### Data analysis

Based on the segmentation of the Haversian canal and the canaliculi, 3D distance transforms were generated. Using the “Exact signed Euclidean Distance Transform (3D)” plugin in FIJI (National Institutes of Health, USA) the calculated 3D distance transforms assign to every voxel of the bone matrix the value of the shortest distance to a compartment surface. The choice of the compartment was either (i) the Haversian canal or (ii) the canaliculi. For all datasets, the distance transforms were computed on the original volumes, but the subsequent correlation analysis was restricted to smaller subvolumes (~25×20×40 μm^3^) to avoid boundary effects. Such acquired distance maps for the canaliculi and the Haversian canal allow to perform a spatial correlation analysis between matrix mineralization (BSE gray value) and the location with respect to a structural compartment (i.e., Haversian canal or canaliculi). The resulting data are presented as gray value probability distribution (GVPD) graphs as a function of the distance to the structural component (see, for example, Fig. 3a). To obtain these distributions, the following steps were performed for each 3D dataset: (i) the gray values (with values ranging from 0 to 255) were plotted versus the distance values to Haversian canal/canaliculi for each voxel using bin sizes of 270 nm/95 nm, respectively; (ii) this initial distribution *m_i, j_* with *m* being the number of voxels with the distance value i and the gray value j was normalized according to

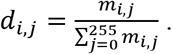

The big advantage of plotting *d_i, j_* instead of *m_i, j_* is that for a fixed value of the distance *i, d_i, j_* represents a gray value probability distribution. Consequently, geometric effects due to a different number of voxels with a given distance are corrected for as the density value is normalized by the sum of all pixels with the same distance. *d_i, j_* denotes the probability of finding a voxel with the gray value *j* at a given distance *i*. Such GVPD plots were used to quantify the mineralization process with respect to the Haversian canal or the canaliculi surface.

## Data availability

The data that support the findings of this study are available from the corresponding author upon reasonable request.

## Code availability

The scripts used during this study are available from the corresponding author on reasonable request.

## Acknowledgements

This study was supported by the AUVA (research funds of the Austrian workers’ compensation board) and WGKK (Vienna Regional Health Insurance Fund).

The authors thank Heike Runge, Birgit Schonert, Gabriele Wienskol, Daniela Gabriel, Sonja Lueger, Petra Keplinger, and Phaedra Messmer for careful sample preparation.

P.F. and R.W. acknowledge the support of the Cluster of Excellence Matters of Activity. Image Space Material funded by the Deutsche Forschungsgemeinschaft (DFG, German Research Foundation) under Germany’s Excellence Strategy – EXC 2025.

## Author contributions

P.F., L.B., and A.R. devised and supervised the study. The samples were prepared by P.R., P.B., and A.B. The staining protocols were designed by M.A., A.R., and L.B. The staining experiments were performed by M.A. FIB/SEM datasets as well as CLSM images were obtained by M.A. Preprocessing of the images were performed by M.A., A.R., and L.B. The scripts for 3D data analysis was written by A.V. Data analysis was conducted by A.V. and M.A. The results were discussed by M.A., A.V, R.W., L.B., A.R., and P.F. The paper and the figures were drafted by M.A., A.V, R.W., and A.R. The manuscript was revised by R.W., L.B., A.R., and P.F., P.R., A.B., P.B. All authors reviewed the paper and commented on it.

## Competing interests

The authors declare no competing interests.

## Correspondence

and requests for materials should be addressed to P.F., A.R., and L.B.

## Notes

### Competing Interest Statement

The authors have declared no competing interest.

