## Supplementary Information for "Three-dimensional interrelationship between osteocyte network and forming mineral during human bone remodeling"

Ayoubi et al.

**Environmental scanning electron microscopy (ESEM).** ESEM imaging (FEI Quanta 600, USA) was performed in order to locate the areas of interest. Using the backscatter signal which provides information on the local average atomic number and therefore on the local mineral content, we searched for forming osteons. Such osteons were identified based on a characteristic spatial gradient in the image brightness corresponding to the transition between unmineralized osteoid and the mineralizing bone like done in a previous study [1]. From the overview images with 50× magnification (2.6 µm pixel size), in each of the three samples, three forming osteons were located and imaged at 600× magnification (270 nm pixel size) under low pressure (1 mbar) with 12.5 kV acceleration voltage and ∼10 mm working distance.

**Confocal laser scanning microscopy (CLSM) of the LCN.** Forming osteons preselected by ESEM were further pre-characterized with an inverted TCS SP8 (Leica, Wetzlar, Germany) CLSM at 40x magnification (HC PL APO 40×/1.30 OIL) to obtain 3D maps of the LCN by mapping the rhodamine signal as previously described [2, 3, 4]. This enables us to define ROIs for FIB-SEM imaging exhibiting an LCN in the mineralized bone/osteoid interface. For excitation, the 514 nm line of an Argon laser was used and the fluorescence signal window was set from 540 nm to 620 nm while using an airy 1 pinhole of 65.4 µm. Depth scans were performed with a typical isotropic voxel size of ~0.38 µm generating 3D datasets of ~300 × 300 × 50 µm³ with a scan speed of 200 lines per second. To compensate for the signal loss with increasing depth, the laser intensity and the photomultiplier gain were gradually increased.

**Selection of ROI for FIB-SEM imaging.** The ROIs for FIB-SEM measurements were selected based on two main criteria. First, to be able to track the course of mineralization, the ROI should include both unmineralized and mineralized tissue and the transition in-between. The ROI should also be entirely located within the forming osteon avoiding adjacent interstitial bone or a neighboring osteon. Second, the ROI should include a clearly distinct part of the lacunocanalicular network (LCN) since one aim of this study was to characterize the interaction of the canaliculi with their surrounding mineralizing matrix. A combination of two imaging techniques, environmental scanning electron microscopy (ESEM) and confocal laser scanning microscopy (CLSM), was used to ensure that these two criteria were met. With ESEM the surface of uncoated samples was imagined discriminating mineralized and unmineralized tissues, and to verify the effectiveness of iodine staining. Supplementary Figure 1a shows the ESEM overview image of sample #1 from which three forming osteons were selected. CLSM was employed to visualize the LCN buried below the sample surface inside the unmineralized and mineralized tissue. For the imaging of the LCN we followed the measurement protocol developed earlier (see details in [2, 3, 4]). By adjusting the laser intensity, the LCN was imaged in different regions of the forming osteon. Using high laser power, the LCN in the fully mineralized matrix could be imaged, whereas the osteoid region produced an oversaturated signal (Supplementary Fig. 1c). Reducing the laser intensity (Supplementary Fig. 1d), the unmineralized region of the osteoid near the Haversian canal could be more clearly investigated, but then the details of the canaliculi network in the mineralized regions were lost. Therefore, both approaches were applied to image the LCN in the mineralized matrix and the osteoid around the Haversian canal. The correlative imaging (ESEM-CLSM), enabled visualization of the local mineral content of the bone tissue with backscattered electron images of ESEM and the LCN with the CLSM in the same region, and the ROI for FIB-SEM was selected accordingly (red rectangle in Supplementary Fig. 1b).

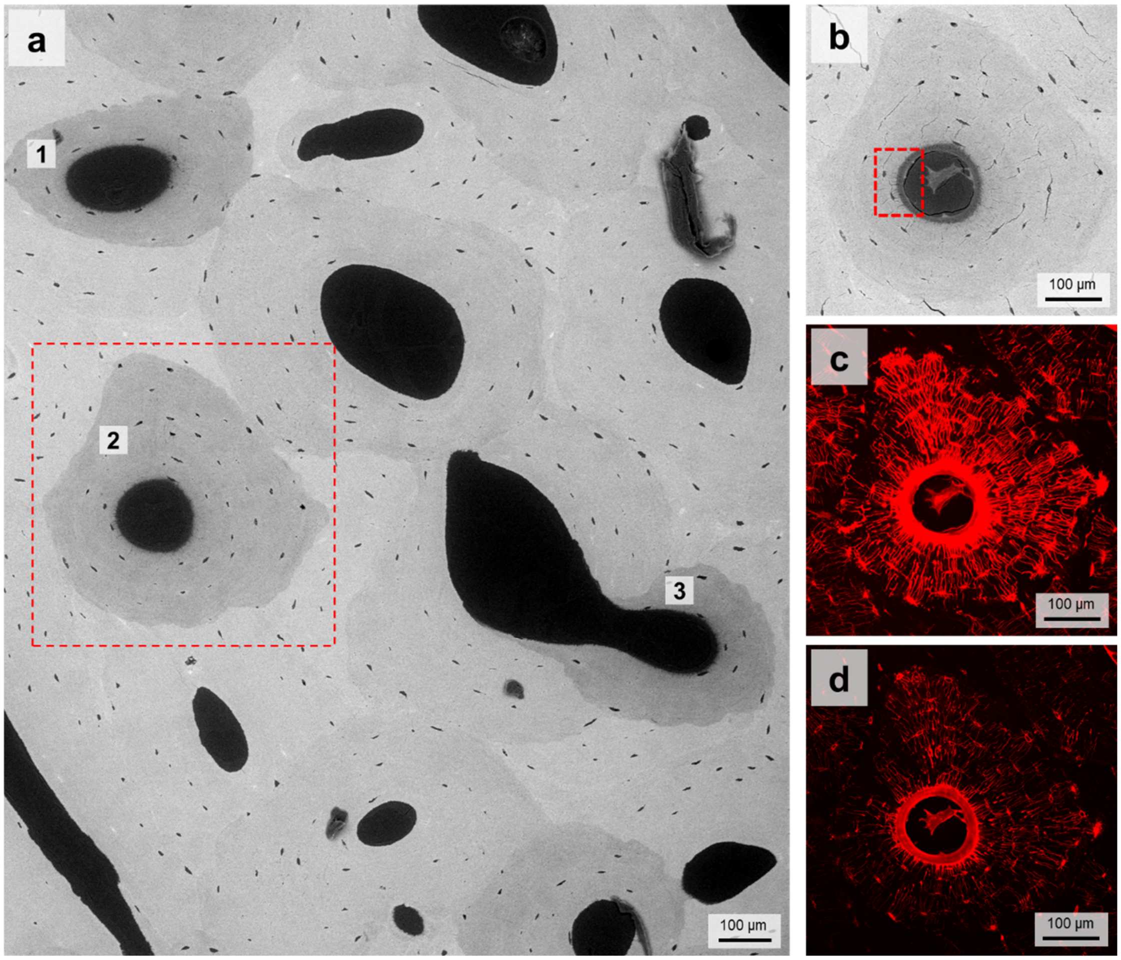


**Supplementary Figure 1. Selection of ROI with ESEM and CLSM for FIB-SEM imaging.** a) Environmental scanning electron microscopy (ESEM) overview of sample #1 from which three forming osteons were selected (dashed square shows osteon #2). b) The ESEM image of osteon #2 after staining, and the ROI (red rectangle). The selected ROI was used for FIB-SEM imaging afterward. c) Confocal laser scanning microscopy (CLSM) image of the same osteon after staining the sample with rhodamine. Note the dense lacunocanalicular network (LCN) within the osteon. The layer thickness for the CLSM dataset was 0.38 µm, and the final depth of imaging was 50 µm. d) The CLSM image with lower laser intensity, showing osteoid and the canaliculi adjacent to it.

**Iodine staining protocol.** For PMMA-embedded bone samples including unmineralized tissue, it is difficult to differentiate between PMMA and soft tissue as both materials consist of mostly light elements such as H, C, N, and O that have small backscattered electron coefficients [5]. To overcome this limitation, soft tissues are often stained with heavy elements-containing chemicals such as uranyl acetate, lead citrate, and osmium tetroxide [6]. Such stains enable to image details of soft tissues in different biological samples. However, in post-embedding staining protocols, such stains might lead to overstaining of polishing blemishes and also mineralized tissue, which is undesirable [7]. Moreover, after embedding, the staining with the chemicals mentioned above is more difficult, because they cannot efficiently penetrate the sample. In contrast, iodine (atomic number Z = 53), is advantageous for staining of archival PMMA-embedded samples without causing the issues mentioned above [7]. The dry protocols rely on gaseous iodine, which is advantageous to avoid problems associated with wetting of specimens, such as specimen shrinkage [8] and pooling around cracks and defects [9] that causes overstaining and consequently higher backscattered signal. In this study, an optimized dry protocol of iodine staining was established for visualization of the osteoid in the forming osteons of PMMA-embedded human osteonal bone. The optimized protocol was conducted in a way that the vapor pressure of iodine was kept below the saturation point of iodine, i.e., < 0.305 mmHg [10]).

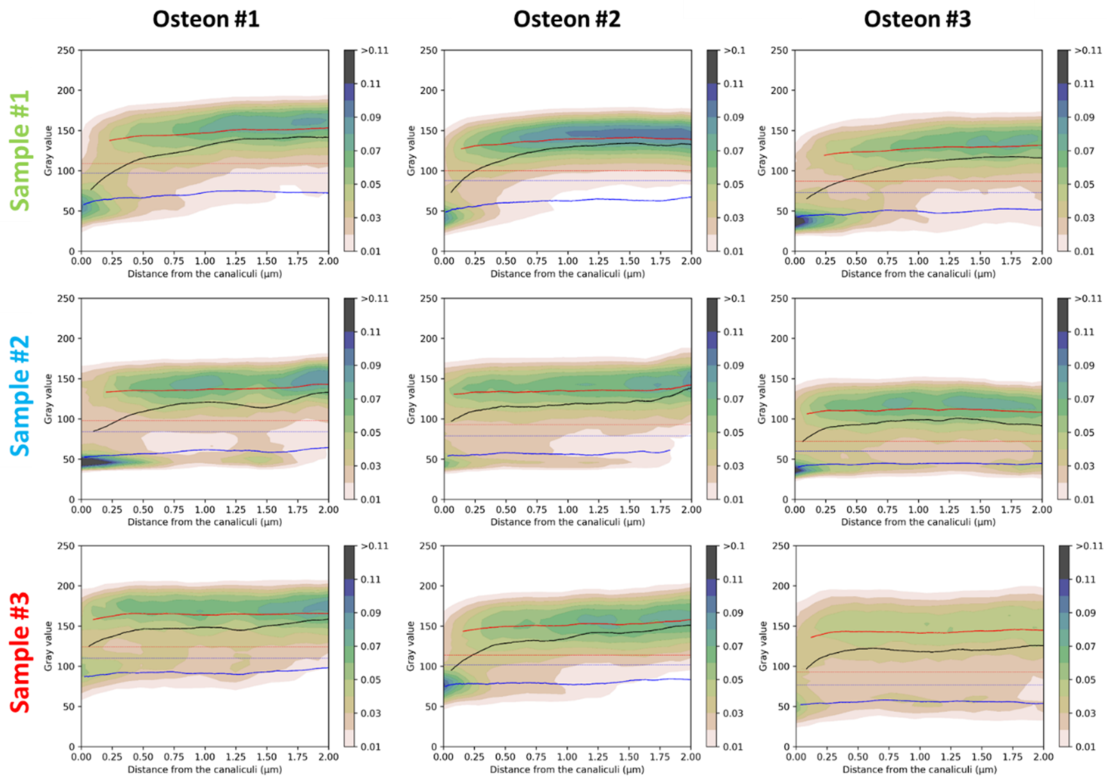


**Supplementary Figure 2. The GVPD graphs of subvolumes with high $V_{MM}/V_{TM}$ for different samples.**
